# Zinc dysregulation in *slc30a8* (*znt8*) mutant zebrafish leads to blindness and disrupts bone mineralisation

**DOI:** 10.1101/2020.09.02.279182

**Authors:** Eirinn W Mackay, Sofía Ibañez Molero, Lavitasha Harjani Tirathdas, Josi Peterson-Maduro, Jingjing Zang, Stephan C.F. Neuhauss, Stefan Schulte-Merker, Stephen W. Wilson

## Abstract

Zinc is an essential cofactor for many cellular processes including gene transcription, insulin secretion and retinal function. Excessive free Zn^2+^ is highly toxic and consequently intracellular zinc is tightly controlled by a system of transporters, metallothioneins (MTs) and storage vesicles. Here we describe the developmental consequences of a missense allele of zinc efflux transporter *slc30a8* (*znt8)* in zebrafish. Homozygous *slc30a8*^*hu1798*^ larvae are virtually blind and develop very little or no bone mineral. We show that zinc is stored in pigmented cells (melanophores) of healthy larvae but in *slc30a8*^*hu1798*^ mutants it instead accumulates in the bone and brain. Supporting a role for pigment cells in zinc homeostasis, *nacre* zebrafish, which lack melanophores, also show disrupted zinc homeostasis. The photoreceptors of *slc30a8*^*hu1798*^ fish are severely depleted while those of *nacre* fish are enriched with zinc. We propose that developing zebrafish utilise pigmented cells as a zinc storage organ, and that Slc30a8 is required for transport of zinc into these cells and into photoreceptors.

## Introduction

Zinc is an essential metal found in all domains of life. Approximately ten percent of the human proteome utilises zinc, including over 1,000 transcription factors (reviewed in Hara et al., 2017). Zinc is particularly critical during development; the offspring of female rats fed a zinc-deficient diet show deformities in the skeleton, brain and eyes (Hurley and Swenert’on, 1966), and in humans, dietary zinc deficiency during gestation is linked to structural birth defects and cognitive impairment (Black, 1998; Uriu‐Adams and Keen, 2010). Intracellular zinc levels are tightly regulated by a system of influx transporters (the ZIP/slc39a family), exporters (the Znt/slc30a family) and sequestering proteins (metallothioneins). Free Zn^2+^ is maintained at a very low concentration since it is a potent inhibitor of mitochondrial respiration, and it is particularly toxic to neurons (Capasso et al., 2005; Dineley et al., 2003).

In mice and humans, fourteen proteins in the Slc39a family transport zinc into the cytoplasm, and nine members of the Slc30a family perform the reverse function (Jeong and Eide, 2013). Slc30a (Znt) family proteins are thought to be Zn^2+^/H+ antiporters (Shusterman et al., 2014). Slc30a1 is ubiquitously expressed on plasma membranes, exporting Zn^2+^ to the extracellular milieu, while Slc30a members 2-9 are localised to internal membranes and direct Zn^2+^ into organelles or vesicles to effect specific functions (Huang and Tepaamorndech, 2013). For example, mice with the *lethal milk* allele (Slc30a4^-/-^) are unable to secrete zinc into breast milk (Huang and Gitschier, 1997), while Slc30a2 fulfils this function in humans (Chowanadisai et al., 2006). Mammalian Slc30a8 is largely restricted to the beta cells of the pancreas where it provides Zn^2+^ to insulin secretory granules. Slc30a8^-/-^ mice are predisposed to developing diabetes but are otherwise healthy (Chimienti et al., 2004; Lemaire et al., 2009). Intracellular zinc is sequestered and delivered to enzymes by metallothioneins (MTs), small redox-sensitive proteins with extraordinarily high affinity for Zn^2+^.

Phylogenetic comparison of the Slc30 family between mammals and teleosts uncovered orthologues of all but mammalian Slc30a3 and Slc30a10 (Feeney et al., 2005). Mammalian Slc30a10 has since been recognized as a manganese transporter (Tuschl et al., 2012), and a fish orthologue identified (Xia et al., 2017). Slc30a3 is the predominant transporter found in the mammalian brain, transporting Zn^2+^ into presynaptic vesicles in glutamatergic neurons (Frederickson et al., 2005). Slc30a3 also maintains zinc homeostasis in the retina (Ugarte and Osborne, 2014). The absence of a direct orthologue of Slc30a3 in teleosts raises the question of whether another member of the Slc30a family provides these important functions. The metallothionein (MT) family is also reduced in teleosts: four mammalian MT genes are known, while two zebrafish MT genes, *mt2* and *mtbl*, have been identified (Chen et al., 2004; Hiu-Mei Yan and Chan, 2002) with *mt2* demonstrating a dose-dependent response to zinc and cadmium (Brun et al., 2014; Wu et al., 2008).

Here we describe the loss of zinc homeostasis and subsequent developmental defects in a zebrafish line (*hu1798*) carrying a mutation in *slc30a8 (znt8)*. Homozygous larvae are almost entirely devoid of mineralised bone, while large deposits of zinc were observed in bone structures and in some cases the brain. Both *slc30a8* and *mt2* are strongly up-regulated in the brain and gut of *slc30a8*^*hu1798*^ larvae suggesting there is a feedback mechanism by which Zn^2+^ levels regulate expression of zinc regulatory genes. In wild-type zebrafish, zinc was found to accumulate in melanophores - pigmented epithelial cells known as melanocytes in mammals - but was absent in these cells in *slc30a8*^*hu1798*^ embryos. *Nacre* zebrafish, which lack melanophores, have accumulations of zinc in the bone and brain but do not show overt defects in mineralisation unless challenged with increased environmental ZnCl_2._

Homozygous *slc30a8*^*hu1798*^ larvae were also found to be completely blind and examination of the retina showed major reductions in the density of photoreceptors, which were also devoid of zinc. Based on these phenotypes and expression patterns, we propose that during development, zebrafish Slc30a8 sequesters zinc in pigmented cells and delivers zinc to photoreceptors.

## Results

### *slc30a8*^*hu1798*^ larvae have reduced bone calcification and dark pigmentation

The *hu1798* mutant was identified in a forward genetic screen for bone defects using alizarin red, a histological stain for calcium (Puchtler et al., 1969; Spoorendonk et al., 2010). Compared to heterozygous and wild-type siblings, homozygous *hu1798* larvae showed substantially less alizarin red staining at 5 days post-fertilisation (dpf) (**Fig**. 1A). The operculum and cleithrum were under-calcified and no staining was visible around the notochord or vertebrae. To help understand the alterations to bone mineral composition in *hu1798* mutants, we utilised the von Kossa stain for phosphate (Rungby et al., 1993). Phosphate was reduced or absent in bone elements of mutants (**Fig**. S1) in a manner consistent with the reduction in alizarin red staining. Together these results indicate a lack of calcium phosphate (hydroxyapatite) mineral in the bone of *hu1798* mutants.

**Figure 1:**
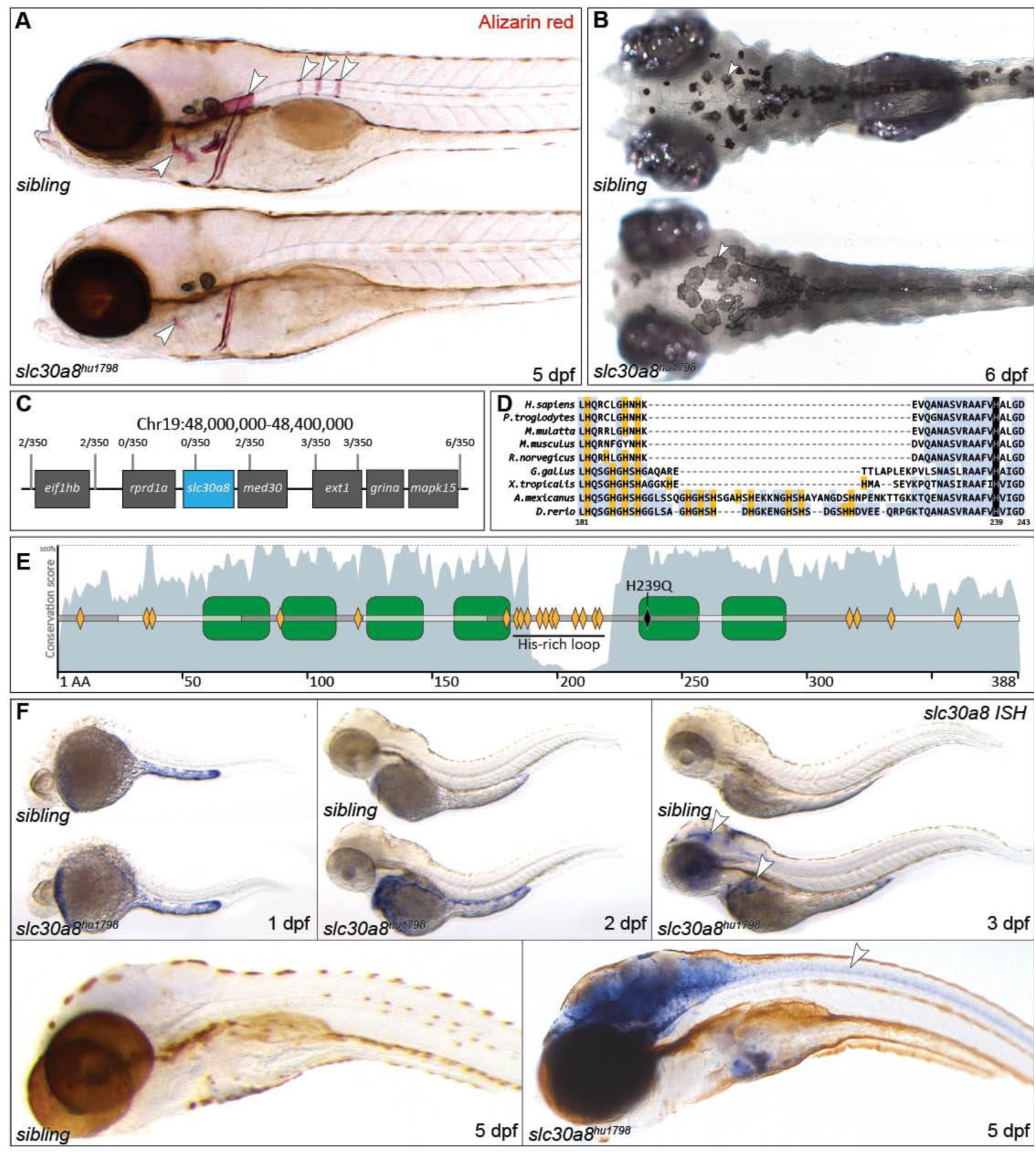
*slc30a8*^*hu1798*^ mutant larvae are characterized by reduced alizarin red staining and increased dorsal pigmentation. **(A)** Alizarin red stain of larvae at 5 days post-fertilisation (dpf) reveals that *slc30a8*^*hu1798*^ bone elements are under-mineralised, particularly the operculum, the anterior notochord and the chordacentra (arrowheads). **(B)** Dorsal view showing enlarged pigment cells in *slc30a8*^*hu1798*^ mutants)(6 dpf). **(C)** Genomic region (assembly version Zv9) linked to the mutation by positional cloning approaches; numbers indicate the frequency of recombinant larvae found at each marker. **(D, E)** The *slc30a8* allele contains a H239Q missense mutation in the 5th transmembrane domain (green boxes). Other histidines are marked orange. ClustalX conservation score for each residue is plotted in grey. Note that some teleosts contain an extra His-rich loop. **(F)** Whole-mount *in situ* hybridization reveals increasing expression of *slc30a8* in the gut, brain, eye and neural tube of the *slc30a8*^*hu1798*^ larva compared to sibling larvae.

Mutant fish did not inflate the swim bladder and were not adult viable. From 5 d.p.f onwards, mutants appeared noticeably darker than siblings due to expansion of pigmented melanosomes across the cytoplasm of the dorsal melanophores (**Fig**. 1B). This change in melanophore morphology is associated with background light adaptation (Logan et al., 2006) and is disrupted when vision is compromised (Neuhauss et al., 1999).

### *hu1798* corresponds to a mutation in *slc30a8 (znt8)*

Using standard positional cloning approaches and bulk segregant analysis, the *hu1798* allele was mapped to a region on chromosome 19 containing two genes, *rprd1a* and *slc30a8* (**Fig**. 1C). The exons of both genes were sequenced revealing an A-T substitution in exon 5 of *slc30a8*. The resulting substitution, H239Q, affects a strictly conserved residue (orthologous to human His 220) in a histidine-rich region which, based on studies in orthologous proteins (discussed below), is predicted to abolish zinc transport (**Fig**. 1D,E). Given Slc30a8 is a zinc transporter and we find defects in zinc localisation in *hu1798* mutants (described below), we conclude that the H239Q mutation in *slc30a8* is causative of the *hu1798* mutant phenotype.

Whole-mount *in situ* hybridization (ISH) revealed expression of *slc30a8* in cells lining the yolk at 1 dpf that declined over time in wild type embryos, whereas in mutants strong expression of the mutant transcript was apparent in the gut, brain, eye and neural tube (**Fig**. 1F). This suggests that compromised Slc30a8 function leads to increased *slc30a8* expression. In support of this, previous studies have found that expression of some zinc transporters can be regulated by changes in intracellular zinc levels via the metal‐responsive element–binding transcription factor‐1 (MTF-1) (Kimura and Kambe, 2016; Laity and Andrews, 2007) and indeed *slc30a8* is up-regulated in the gills of adult zebrafish when excess zinc is present in the water (Feeney et al., 2005). Despite the bone phenotype in *hu1798* mutants, expression of *slc30a8* was not detectable by ISH in cells associated with mineralized elements.

### Zinc is abnormally distributed in *slc30a8* mutants

Mammalian pancreatic beta cells can be identified using the histological stain dithizone which produces an orange precipitate upon reaction with Zn^2+^ (Danscher et al., 1985), but the beta cells of *slc30a8*-null mice are negative for this stain as they contain insufficient zinc (Lemaire et al., 2009). We applied the dithizone stain to larvae to determine if zinc was altered in the pancreas of mutants. While staining was not detected in the pancreas of wild type or mutant larvae, the stain was visible in the head and craniofacial bone elements of *slc30a8*^*hu1798*^ larvae from 4 d.p.f onwards, while siblings showed little or no staining anywhere (**Fig**. 2A). This suggests that the loss of calcium phosphate is accompanied by increased zinc in forming bones. Supporting this, the fluorescent sensor, TSQ (6-methoxy-8-p-toluenesulfonamido-quinoline) which forms TSQ-Zn-protein adducts (Frederickson et al., 1987; Meeusen et al., 2011) revealed zinc in bone elements, brain and neural tube in *slc30a8*^*hu1798*^ larvae but not in siblings (**Fig**. 2B). Conversely, in siblings TSQ-Zn fluorescence revealed zinc deposits in the skin in a pattern resembling melanophores, while no such fluorescence was visible in *slc30a8*^*hu1798*^ larvae (**Fig**. 2C).

**Figure 2:**
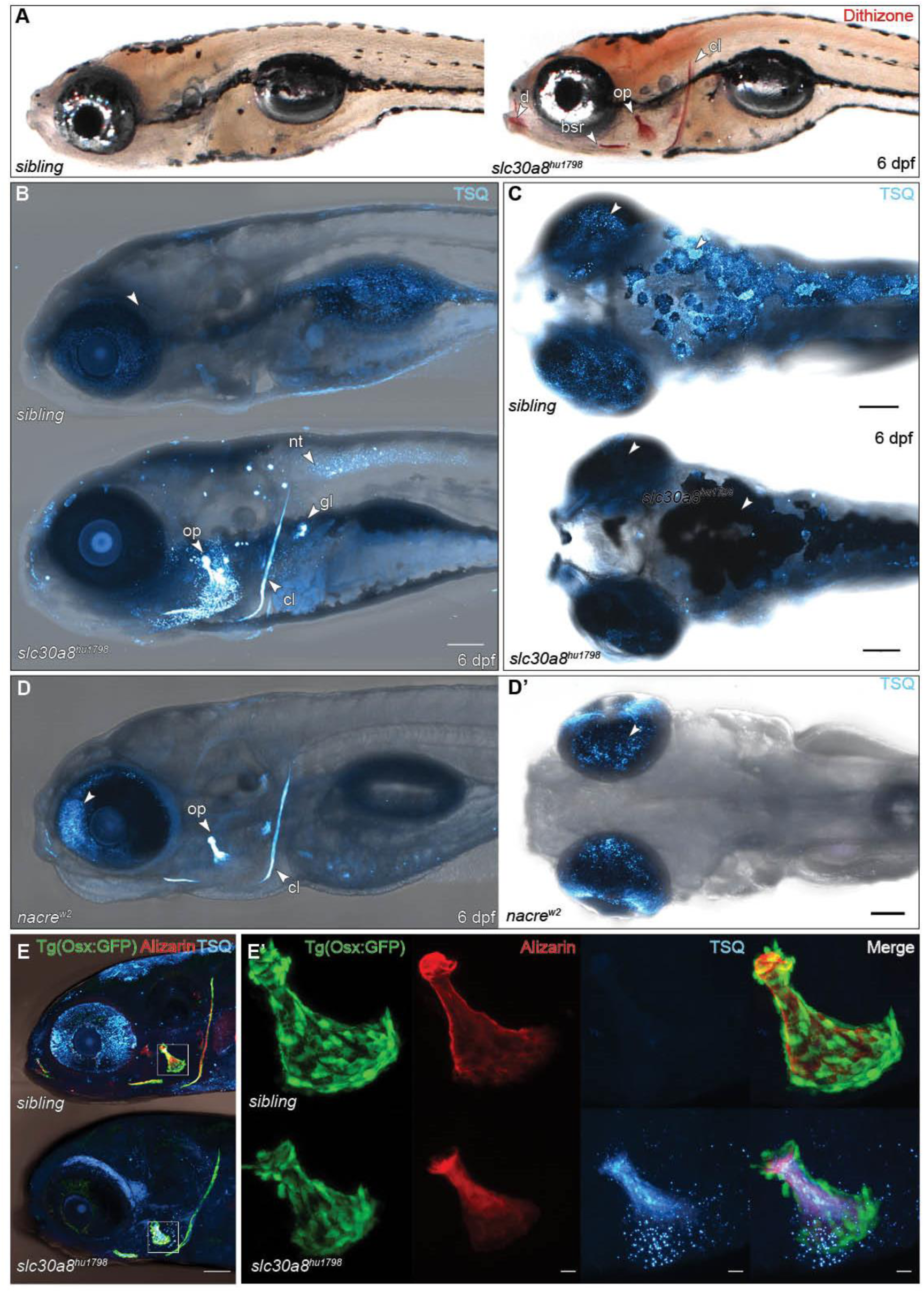
Zinc distribution is altered in *slc30a8*^*hu1798*^ larvae. **(A)** Dithizone stain reveals zinc to be concentrated in craniofacial bone elements such as the cleithrum (cl), operculum (op), branchiostegal ray (bsr) and dentary (d), and the brain of the *slc30a8*^*hu1798*^ larva. **(B)** TSQ fluorescence confirms increased zinc in bone elements of mutant larvae compared to those of siblings. Fluorescence was also observed in the neural tube (nt) and in kidney glomeruli (gl). **(C)** Dorsal view showing TSQ-Zn is normally associated with melanophores and in the eye, but this pattern is absent in *slc30a8*^*hu1798*^ larvae. **(D)** Lateral and dorsal views of *nacre*^*w2*^ larvae, which lack melanophores, showing zinc accumulation in bones and eyes. (E) Mutant larvae still contain active osteoblasts as revealed by expression of the marker *osterix*:GFP around the operculum. **(E’)** Magnification of the operculum of 6 dpf larvae. In *slc30a8*^*hu1798*^ larvae this bone element is smaller, with less mineral as shown by alizarin staining. TSQ staining shows that zinc is associated with the bone mineral and not the *osterix*-positive osteoblasts. Scale bar in (B-D) = 100 µm, (E) = 10 µm.

These results suggest that the normal accumulation of zinc in melanophores fails in *slc30a8* mutants and this leads to accumulation in bone and other sites. To explore this idea, we examined zinc distribution in *nacre*^*w2*^ larvae, which lack melanophores due to a mutation in the *mitfa* transcription factor required for differentiation of melanophores from the neural crest lineage (Lister et al., 1999). TSQ-Zn fluorescence was absent in the skin of *nacre*^*w2*^ larvae whereas, similar to *slc30a8*^*hu1798*^ larvae, zinc accumulated in the bone (**Fig**. 2D). Compared to siblings, the eyes of *nacre*^*w2*^ larvae displayed more intense TSQ-Zn fluorescence while none was visible in the eyes of *slc30a8*^*hu1798*^ larvae.

To examine the relationship between zinc and the under-calcified bone phenotype of *slc30a8*^*hu1798*^ larvae, we assessed expression of the osteoblast marker *osterix:GFP* (Spoorendonk et al., 2008) and stained larvae with both TSQ and alizarin red. GFP was clearly visible in osteoblasts adjacent to mineralising craniofacial elements such as the operculum and cleithrum (**Fig**. 2E). The operculum was smaller in mutant larvae and alizarin red staining was diminished and poorly defined, while TSQ stained the mineral matrix and not the osteoblasts themselves. This indicates that zinc is interacting with the bone mineral directly, and therefore the reduction in calcium hydroxyapatite is likely to be due to excess Zn^2+^ ions in the mineral, rather than through inhibition of osteoblast function by Zn^2+^.

### Bone mineralisation in *nacre* fish is affected by zinc

Unlike *slc30a8*^*hu1798*^ fish, which are not adult-viable, *nacre* fish are healthy, and juveniles stained with TSQ revealed a striking zinc-rich skeleton, while siblings predominantly showed TSQ-Zn in melanophores (**Fig**. 3A). Alizarin red staining did not reveal a difference in bone mineralisation of untreated *nacre* larvae, but the addition of 10 µM ZnCl_2_ to the embryo media from 6 h.p.f to 5 d.p.f resulted in *nacre* larvae with bone elements completely devoid of calcium hydroxyapatite while a moderate reduction was observed in siblings (**Fig**. 3B). At a higher dose of ZnCl_2_ (100 µM) mineralisation was inhibited regardless of genotype (**Fig**. 3C). Both 10 µM and 100 µM concentrations proved lethal to *slc30a8*^*hu1798*^ embryos, most of which died before emerging from the chorion. These results give weight to the notion of Zn^2+^ as a mineralisation inhibitor and suggest the altered zinc homeostasis detected in *nacre*^*w2*^ fish is a milder form of the *slc30a8*^*hu1798*^ phenotype, perhaps due to the eyes of nacre larvae acting as a secondary reservoir for zinc (discussed below).

**Figure 3:**
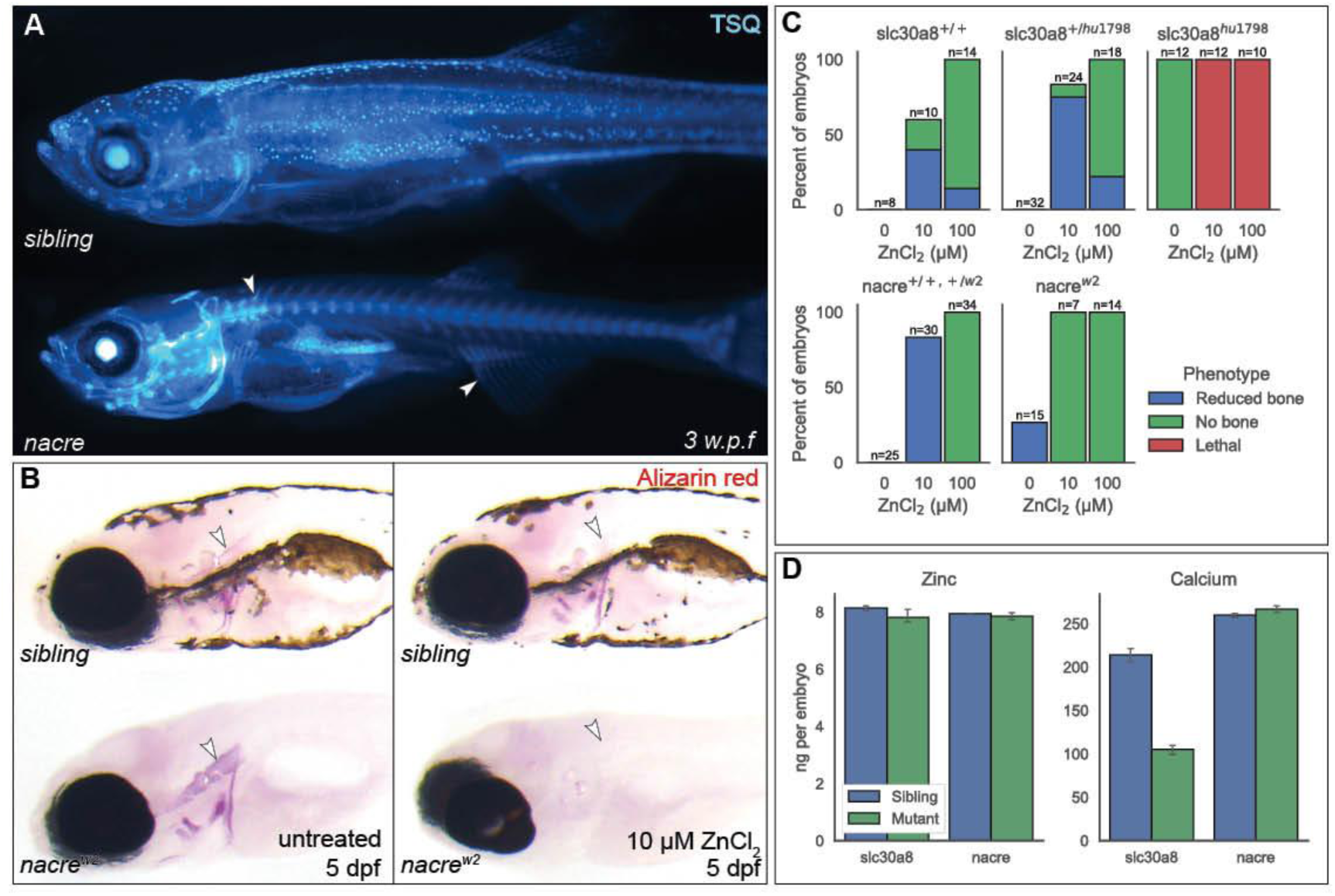
Bone mineralisation in *nacre* fish is affected by zinc. **(A)** Widefield fluorescent image of *nacre*^*w2*^ and sibling juveniles at 3 weeks’ post-fertilisation demonstrating TSQ fluorescence in the skeleton of the *nacre* fish, notably in the spine and fins. **(B)** Alizarin red bone stains of *nacre* and sibling larvae incubated with or without 10 µM ZnCl_2_ in the media. Arrowheads indicate the notochord. **(C)** Summary of phenotypes generated by ZnCl_2_ treatment from 1 dpf to 5 dpf in *nacre* and *slc30a8*^*hu1798*^larvae. **(D)** Total zinc and calcium content per embryo as determined by ICP-AES. Columns show mean values ±SEM across 3 groups of pooled larvae.

Neither *slc30a8*^*hu1798*^ nor *nacre* larvae showed a difference in total Zn content as measured by inductively coupled plasma atomic emission spectroscopy (ICP-AES), while Ca content was 2-fold lower in *slc30a8*^*hu1798*^ larvae reflecting the under-mineralised bone (**Fig**. 3D). This suggests that the phenotypic defects in *slc30a8*^*hu1798*^ and *nacre* embryos result from altered distribution rather than altered overall levels of zinc.

### Metallothionein levels are up-regulated in *slc30a8*^*hu1798*^ larvae

As MT proteins help to maintain Zn homeostasis by sequestering excess zinc, we speculated that expression of the zebrafish intracellular zinc-sequestering protein metallothionein (MT) gene *mt2* may be altered in *slc30a8*^*hu1798*^ fish as a response to altered zinc distribution. To assess *mt2* levels we detected the endogenous *mt2* gene by ISH and also generated a transgenic line using a fragment of the *mt2* promoter, previously shown to be zinc-responsive (Chen et al., 2004), upstream of a cassette encoding GFP.

High levels of *mt2* expression were evident in the brain, eyes, neural tube and gut of 3 dpf *slc30a8*^*hu1798*^ larvae but undetectable in siblings (**Fig**. 4A). GFP fluorescence was observed in the same tissues as detected by ISH, while in siblings GFP was restricted to a few isolated cells in the skin (**Fig**. 4B). GFP was most strongly expressed in the brain from 4 dpf onwards, preceding the appearance of TSQ-Zn in the brain at 6 dpf. In both wild type and *slc30a8*^*hu1798*^ larvae and across all tissues, GFP and TSQ-Zn occupied distinct territories and were not observed in the same cells (**Figs**. 4B-D). This spatial separation between *mt2:gfp* and TSQ-Zn is most clearly evident in the neural tube (**Fig**. 4D), where GFP-positive cells resembling neurons were adjacent to (but not co-located with) many smaller (subcellular) TSQ-Zn bodies. Drawing on a study reporting that (under normal conditions) zebrafish *mt2* expression in the brain is localised to neurons and not glial cells (Teoh et al., 2015), these observations suggest that distinct neural cell populations respond to disrupted zinc homeostasis in different ways, with neurons increasing MT activity and others developing zinc inclusions.

**Figure 4:**
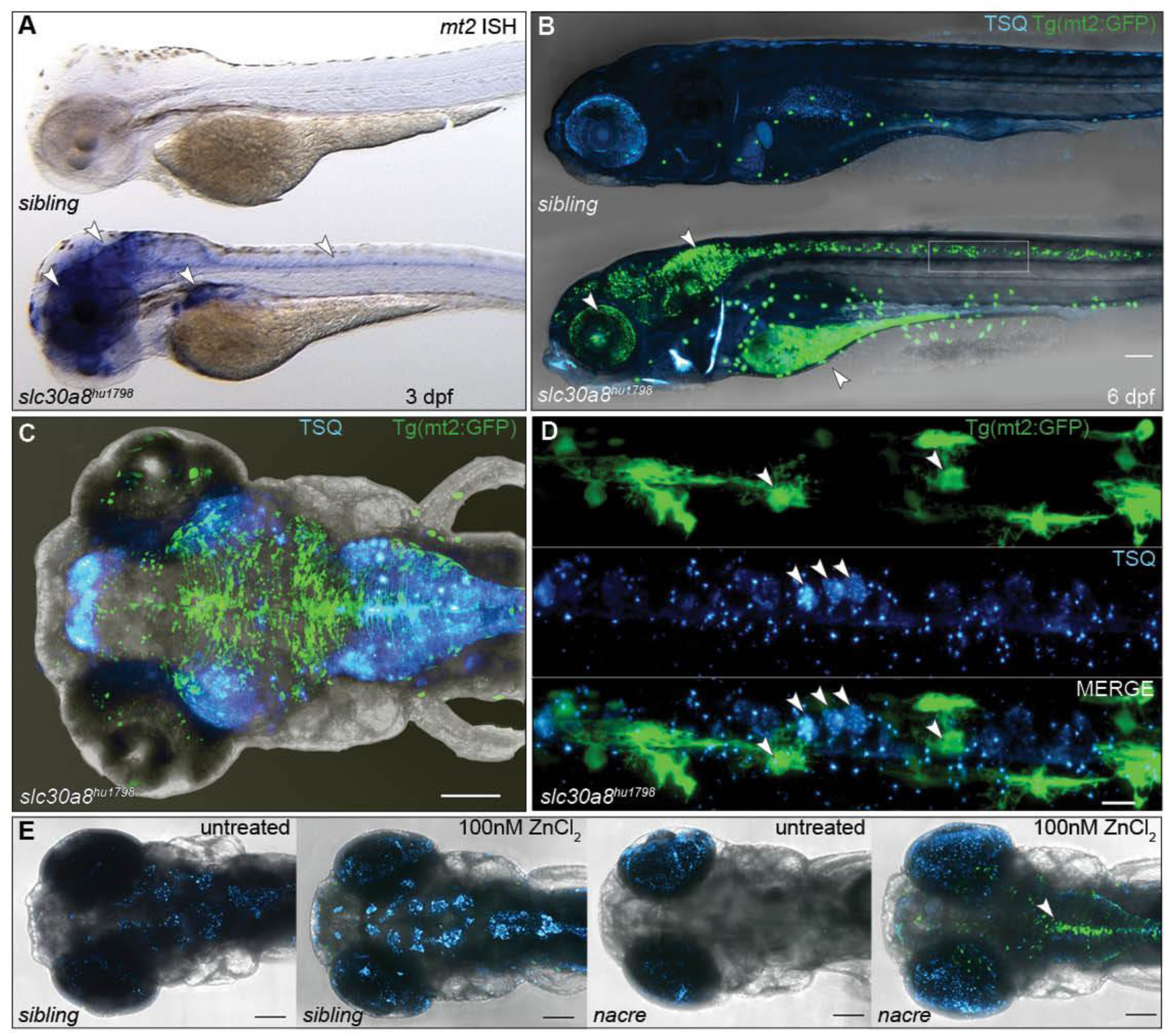
Metallothionein (*mt2*) expression is up regulated in *slc30a8*^*hu1798*^ larvae. **(A)** Whole mount ISH showing extremely elevated expression of *mt2* in the brain, eye, gut and neural tube of a *slc30a8*^*hu1798*^ larva. **(B)** A transgenic reporter *mt2:gfp* is expressed in the same tissues demonstrated in (A), and additionally in the gut. The box indicates the neural tube region magnified in (D). **(C)** Dorsal view of a *slc30a8*^*hu1798*^ larva showing *mt2:gfp* expression and TSQ-Zn fluorescence. **(D)** Magnification of the neural tube of a *slc30a8*^*hu1798*^ larva, showing spatial separation between cells expressing *mt2:gfp* and cells containing deposits of TSQ-Zn. (E) TSQ fluorescence and *mt2:gfp* expression in sibling and *nacre*^*w2*^ larvae incubated with low-dose (100 nM) ZnCl_2_ from 4 to 6 dpf. Scale bar in (B, C, E) = 100 µm, (D) = 10 µm.

To help validate the notion of *nacre*^*w2*^ larvae featuring a mild form of zinc dysregulation, we examined *nacre*^*w2*^ larvae expressing *mt2:gfp* which revealed GFP fluorescence in a pattern identical to siblings (i.e, none or very few cells expressing GFP). Following incubation with ZnCl_2_ concentrations as low as 0.1 µM, however, distinctive GFP expression was observed in the hindbrain and neural tube (**Fig**. 4E). Sibling larvae under these conditions simply showed an increase in zinc content of melanophores.

### Visual function is severely impaired in *slc30a8* mutants

The reduced levels of zinc in the eyes of *slc30a8*^*hu1798*^ larvae, together with the increased pigmentation (suggesting a failure of background adaptation) led us to consider that there might be some sort of visual impairment in *slc30a8*^*hu1798*^ larvae, resulting in blindness. We tested visual function by measuring the optokinetic response (OKR) in 6 dpf larvae. This experiment found oculomotor movements in *slc30a8*^*hu1798*^ mutants to be nearly absent under a range of contrasts, spatial or temporal frequencies (**Fig**. 5A, repeated-measures ANOVA, P< 0.001). In order to directly test the involvement of the outer retina in this visual defect, we performed the electroretinogram (ERG) to measure the field potential of the retina. **Fig**. 5B presents ERG traces recorded from sibling and mutant larvae. The b-wave amplitude is proportional to stimulus intensity before saturation and hence a reliable read-out of outer retina function (**Fig**. 5C). In mutants, although the b-wave amplitude was still intensity dependent, the amplitude was dramatically reduced (repeated-measures ANOVA, P<0.001) compared to siblings. Since the b-wave amplitude was reduced by more than 90% in *slc30a8*^*hu1798*^ larvae and there was still no measurable a-wave, the defect is likely connected to light perception in photoreceptors.

**Figure 5:**
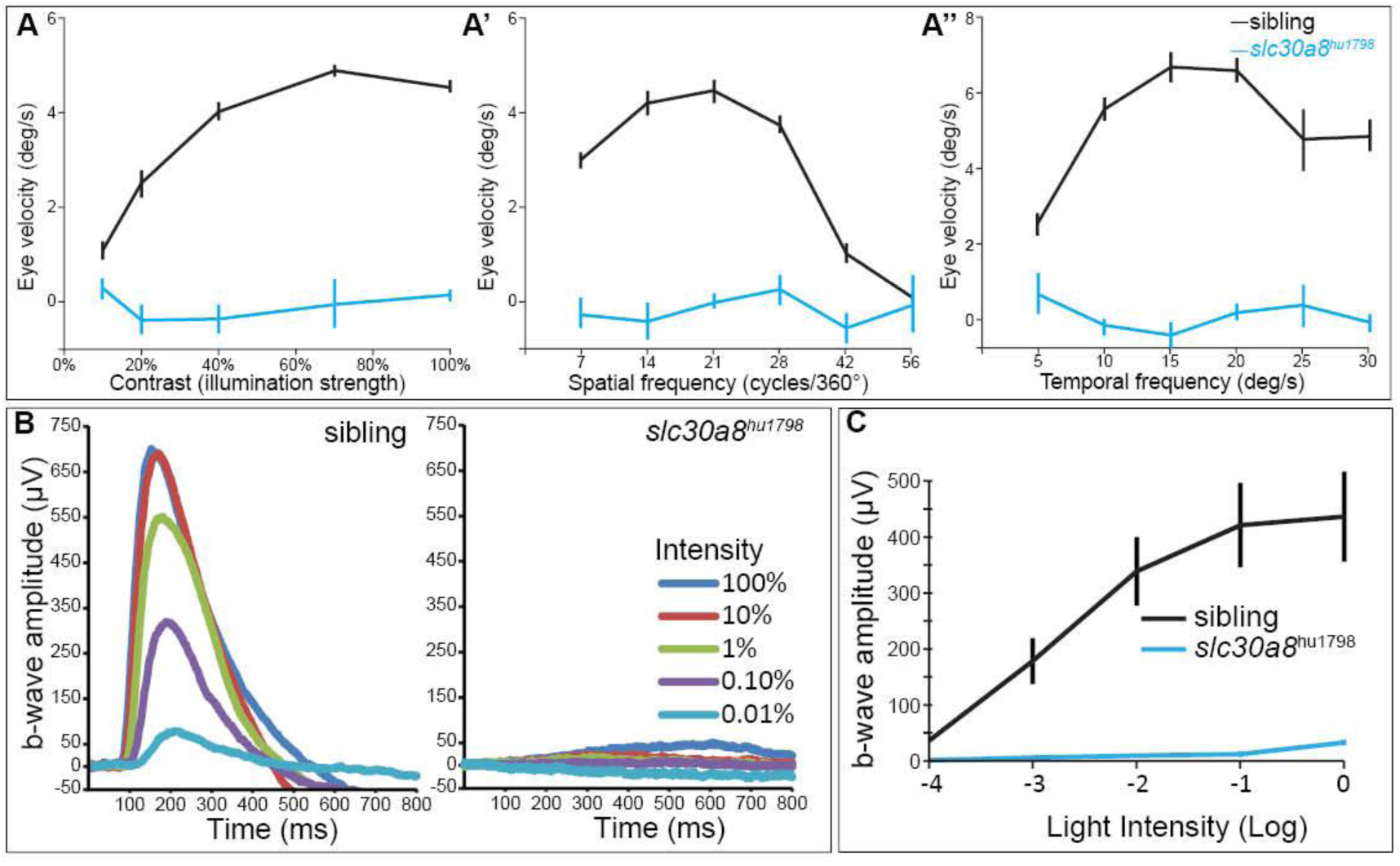
*Slc30a8*^*hu198*^ mutant larvae exhibit visual impairments. **(A)** Eye velocity elicited by the optokinetic response (OKR) was measured at different contrast (5%, 10%, 20%, 40%, 70% and 100% of full illumination) with a spatial frequency of 20 cycles/360° and an angular velocity of 7.5 deg/s. **(A’)** Eye velocity measured at different spatial frequency (7, 14, 21, 28, 42, 56 cycles/360°) with contrast of 70% and an angular velocity of 7.5 deg/s. **(A’’)** Eye velocity measured at different temporal frequency (5, 10, 15, 20, 25, 30 deg/s) with contrast of 100% and a spatial frequency of 20 cycles/360°. **(B)** Sample electroretinogram (ERG) recordings of a sibling and *slc30a8*^*hu1798*^larvae (6 dpf) in response to flashes at different light intensity (100%, 10%, 1%, 0.1% and 0.01% of maximum intensity). Flash duration was 100 ms and the interval was 10 s. Each trace is the average of two responses. **(C)** Averaged data collected from sibling and *slc30a8*^*hu1798*^ larvae (n=5 each) showing the dependence of ERG b-wave amplitude on the relative light intensity.

### Evidence of photodamage in the retina of *slc30a8* mutants

To assess zinc distribution in the retina we performed TSQ staining on plastic sections. Due to strong autofluorescence in fixed eye tissue (particularly the photoreceptors), images were taken before and after TSQ incubation and the difference calculated using software (ImageJ). SYTOX Green was used to label nuclei and BODPIY-TR methyl ester was used to label the photoreceptor outer segments (OS) (Cooper et al., 2005). In siblings, intense TSQ-Zn fluorescence was observed in the photoreceptor OS, while in *slc30a8*^*hu1798*^ larvae the OS layer was not visible at all, and the outer plexiform and nuclear layers (OPL, ONL) were disorganised. TSQ was visible in the RPE of both genotypes. (**Fig**. 6A). Based on these observations we infer that the TSQ-Zn fluorescence we observed in the eyes of whole-mount siblings but not mutants (Fig. 2C) is emanating from the photoreceptors. To test this, we examined the retinas of sibling and *nacre*^*w2*^ larvae under normal conditions or after treatment with 50 µM ZnCl_2_. As expected, TSQ-Zn fluorescence in the photoreceptors became more intense after this treatment, and particularly so in the *nacre*^*w2*^ larvae (**Fig**. 6B), echoing the observations in Fig. 4E. The photoreceptors of zinc-treated nacre larvae did not appear to be damaged suggesting that functional Slc30a8 allows retinas to tolerate elevated zinc content.

**Figure 6:**
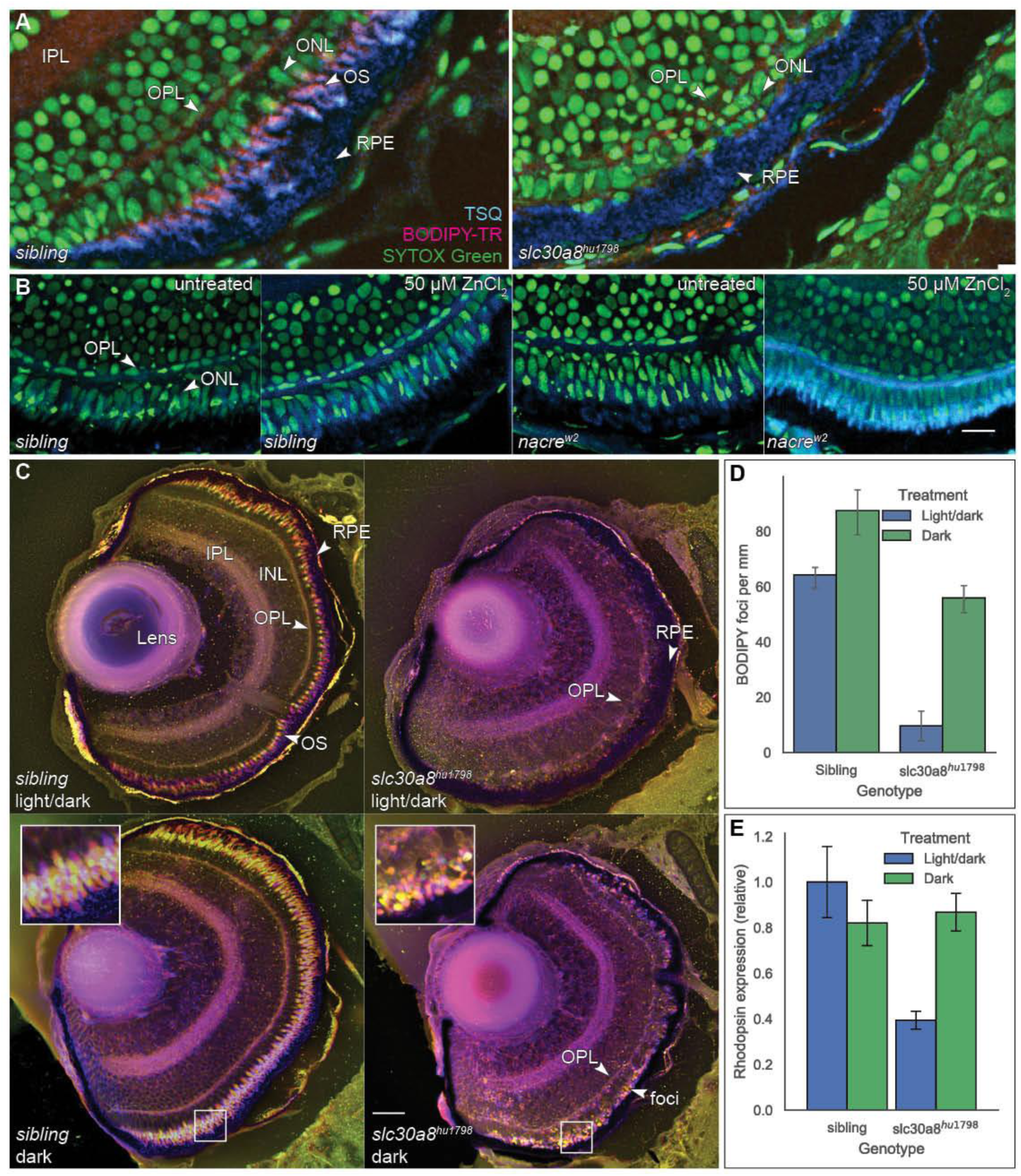
The eyes of mutant larvae exhibit signs of photodamage caused by loss of zinc distribution. **(A)** Sections of the eye stained with SYTOX Green to label nuclei, BODIPY-TR methyl ester to label photoreceptor outer segments (OS) and TSQ to label zinc. OPL: outer plexiform layer. IPL: inner plexiform layer. ONL: outer nuclear layer. RPE: retinal pigment epithelium. **(B)** *nacre*^*w2*^ larvae treated with ZnCl_2_ showed an increase in TSQ-Zn fluorescence in the photoreceptor OS. **(C)** Histological sections from the eyes of 6 dpf larvae raised in normal light/dark cycles or complete darkness, stained by BODIPY-TR and displayed with depth-colour-coding to highlight the photoreceptor OS. White squares indicate the magnified (inset) views. **(D)** Density of BODIPY-positive foci in retinal outer segments (n=4 per group). **(E)** Expression of rhodopsin as measured by qPCR and normalised to the sibling light/dark sample (n=3 per group). Scale bar in (A, B) = 10 µm, (C) = 100 µm.

As zinc is known to play a protective role in the retina (Ugarte and Osborne, 2014) we tested the hypothesis that mutant photoreceptors might be damaged by light by raising embryos in complete darkness (or a normal light/dark cycle) to 6 dpf and staining plastic sections with BODIPY-TR methyl ester. Compared to siblings, the retinas of mutant larvae raised under normal lighting were virtually devoid of BODIPY-positive foci (**Fig**. 6C). Raising *slc30a8*^*hu1798*^ embryos in darkness led to an increase in the number of BODIPY-positive foci (**Fig**. 6C,D; mean 73 (SEM 3.2) VS 12 (SEM 2.69) cells per mm; n=4); however these foci were small and irregular compared to the organised pattern of photoreceptors in wild-type retina. The outer plexiform layer, normally highly disrupted in mutants, was also partially restored in embryos raised in darkness. Supporting the BODIPY labelling data, quantitative PCR (qPCR) analysis of rhodopsin expression suggested an increase in rod photoreceptor mRNA when mutant embryos were reared in the dark (**Fig**. 6E; expression relative to sibling: 0.39 (SEM 0.04) VS 0.86 (SEM 0.08); n=3). Although dark rearing improved retinal structure and opsin expression levels, OKR experiments on dark-raised mutant larvae failed to elicit a response, indicating that the larvae still have a visual deficiency (data not shown). These results suggest that photoreceptor loss in *slc30a8*^*hu1798*^ mutants is partially mediated by light but raising larvae in darkness is not sufficient to restore visual function.

## Discussion

In this study, we have shown that as development proceeds, wild-type zebrafish larvae accumulate zinc in melanophores and photoreceptors, while *slc30a8*^*hu1798*^ larvae do not and consequently accumulate zinc in bone elements and the brain instead. Furthermore, mutant larvae are blind and almost devoid of bone mineral. We draw three conclusions from these results: a major (perhaps primary) role of Slc30a8 is to deliver zinc into melanophores and photoreceptors; photoreceptors die without zinc; and excessive zinc inhibits bone mineralization.

### Function of Slc30a8

The mutation identified in *slc30a8*^*hu1798*^ larvae of a strictly-conserved residue, H239Q, is expected to abolish the function of the Slc30a8 (Znt8) transporter; the bacterial counterpart of this particular residue (H153 of *E. coli* YiiP) directly binds to Zn^2+^ as observed by X-ray crystallography, and substitutions of it abolish zinc transport (Lu and Fu, 2007). In the mutant phenotype described here, this loss of transport results in a loss of zinc in melanophores but the exact mechanism underlying this phenotype remains elusive. Mammalian Slc30a8 transports zinc out of the cytoplasm into insulin vesicles; perhaps zebrafish Slc30a8 is similarly located on the membrane of melanosomes, where it supplies zinc to be stored within the melanin pigment. Using ISH we were unable to detect *slc30a8* expression in melanophores using *tyr* as a positive marker (**Fig**. S2) but could detect it in sections of RPE of mutants (**Fig**. S3). A previous study using RNAseq reported an enrichment in *slc30a8* transcripts in both melanophores and the RPE (Higdon et al., 2013) compared to whole larvae at 3 dpf.

The association between zinc and pigmented tissues has been noted from studies of the mammalian iris (Bowness et al., 1952; Kokkinou et al., 2004), and an affinity of melanin for inorganic ions such as Zn^2+^ and Cu^2+^ has been reported from *in vitro* studies (Potts and Au, 1976; Sarna et al., 1980). More recently, X-ray absorption spectroscopy analysis of pigmented bird feathers showed the distribution of zinc (along with copper and calcium) to be highly correlated with the distribution of melanin (Edwards et al., 2016). It seems this affinity is exploited - at least in developing zebrafish - to provide a zinc sequestration pool. An examination of the sub-cellular location of Slc30a8 will yield insights into this sequestration process.

### Zinc as a mineralisation inhibitor

Is the hypo-mineralised phenotype of *slc30a8*^*hu1798*^ and *nacre* larvae (when challenged with Zn^2+^) directly caused by excessive zinc in the bone itself? Zinc is a normal component of bone, and zinc deficiency is associated with low bone mass in rats and humans (Eberle et al., 1999; Hyun et al., 2004). Zinc supplementation also increases bone density in rats and healthy humans (Seco et al., 1998; Peretz et al., 2001), particularly in patients with low zinc status (Fung et al., 2013). Studies in vitro, however, have shown that Zn2+ ions inhibit the formation of calcium hydroxyapatite crystals, a notion which we here confirm in vivo; the Zn^2+^ nucleus is larger than Ca^2+^, so its inclusion distorts the growing crystal lattice (“crystal poison”) (Bigi et al., 1995; Kanzaki et al., 2000; Chaikina et al., 2020). Zinc-doped hydroxyapatite is of great interest as a biomaterial as it appears that low concentrations of zinc (0.3-1.6% w/w) appears to enhance bone regeneration when used as an implant (or implant coating) compared to HA alone (Kawamura et al., 2000; Tao et al., 2016; Thian et al., 2013) via several mechanisms including inhibition of bone resorption (reviewed in Cruz et al., 2018).

Zinc may be beneficial for mature bone mineral density in humans, but it appears that zinc accumulation to the degree observed in *slc30a8*^*hu1798*^ larvae is pathological for bone formation. Further insights into this balance may come from exploiting the propensity of *nacre* larvae to take up ZnCl_2_ from the media into the bone elements.

### Zinc deficiency in the eye causes visual impairment

In our experiments in *slc30a8*^*hu1798*^ larvae, we observed missing photoreceptor outer segments and disrupted outer plexiform and nuclear layers, a 2-fold reduction in rhodopsin expression, severely reduced ERG b-waves, and eyes which cannot follow moving targets. The morphological disruption in the retina is exacerbated by exposure to light. We speculate that this is the consequence of a lack of zinc transport from the RPE to the photoreceptors. An alternative hypothesis is that, similar to the bone phenotype, retinal degeneration in *slc30a8*^*hu1798*^ larvae may be caused by toxic levels of zinc resulting from the loss of melanophore-provided sequestration. We treated *nacre*^*w2*^ larvae with ZnCl_2_ at a concentration which inhibited bone mineralisation and did not observe overt disruption of photoreceptor morphology; in fact, photoreceptors appeared to readily take up zinc in the outer segments. Presumably, then, Slc30a8 has another role in the retina, unrelated to zinc sequestration in the melanophores. The RPE is known to contain a high concentration of zinc and most ZIP and Znt transporters (including Slc30a8) are present in these cells (Leung et al., 2008). We propose that Slc30a8 facilitates the transport of zinc from the RPE to the photoreceptors.

Zebrafish, as other vertebrates, have two functional visual cycles, the canonical involving the retinal pigment epithelium (RPE) and a cone-specific alternative cycle involving Muller glia cells (reviewed in Fleisch and Neuhauss, 2010). Mutations in most genes coding for components of both pathways are linked to outer retinal dystrophies (reviewed in Berger et al., 2010), providing a rationale for the observed phenotype in *slc30a8* mutants. Zinc influences the function of a number of proteins in the visual transduction cascade (Ugarte and Osborne, 2014), such as the recycling of all-*trans* retinol into 11-*cis* retinal by the zinc-dependent enzyme, retinal dehydrogenase. Mice carrying a null mutation for this enzyme experience light-mediated apoptosis of the photoreceptors (Maeda et al., 2006). Zinc deficiency is also linked to caspase-dependent apoptosis of cultured retinal cells, including RPE, photoreceptor and retinal ganglia cells (Hyun et al., 2000; Shindler et al., 2000; Tamada et al., 2007). In rats kept on a zinc-deficit diet for several weeks, rod photoreceptor outer segment degeneration has been reported (Leure-duPree and McClain, 1982). In humans, zinc deficiency has been shown to lead to defects in the scotopic (dark adapted) ERG (Mochizuki et al., 2006) indicating a prominent role of zinc for rod function.

## Conclusion

We note two distinct functions of Slc30a8 in zebrafish. Our results show that Slc30a8 is required for storing zinc in melanophores which appear to function as a major zinc reservoir. The loss of this function during development leads to accumulation of zinc in other tissues (notably the brain and bone elements) with developmental/pathological consequences. The second function is in the eye, where we propose Slc30a8 maintains retinal health by transporting zinc to photoreceptors from the RPE. In the case of *nacre* fish which lack melanophores, melanophore zinc storage is unavailable but the RPE is intact and zinc is still delivered to the photoreceptors. This allows for normal vision and apparently provides a secondary zinc reservoir which may explain why the *nacre* phenotypes are mild compared to that of slc30a8^*hu1798*^ mutants. The discovery here of two zebrafish lines with altered zinc homeostasis provides an opportunity to investigate the role of zinc as an inhibitor (and perhaps, an enhancer) of bone mineralisation as well as its effect on retinal function, using a highly tractable model organism.

## Materials and Methods

### Zebrafish husbandry and positional cloning

Zebrafish were maintained under standard husbandry conditions according to FELASA guidelines (Aleström et al., 2019). Larvae were raised in E3 media (5 mM NaCl, 17 µM KCl, 330 µM CaCl2, 330 µM MgSO4) at 28°C. A forward genetic screen using ethylnitrosourea (ENU)-induced mutagenesis and alizarin red to examine bone development was performed as described previously (Huitema et al., 2012; Spoorendonk et al., 2010). The mutation in *slc30a8* was mapped using positional cloning (Geisler, 2002) and confirmed by Sanger-sequencing. Subsequent genotyping of the *hu1798* allele was performed using the KASP assay mix (LGC Genomics, Hoddesdon UK) with the following primers:

Wildtype: 5’-GAAGGTGACCAAGTTCATGCTCAGCAGATCTCCAATCACA-3’,

Mutant: 5’-GAAGGTCGGAGTCAACGGATTAGCAGATCTCCAATCACT-3’,

Common reverse: 5’-GCTAGTGTCCGGGCGGCGTT-3’.

### Whole-mount in situ hybridization

ISH was performed as described (Schulte-Merker, 2002; Thisse and Thisse, 2008). The templates for *slc30a8, mt2* and *tyr* were amplified from zebrafish cDNA using the following primer pairs. Probes were detected with HRP anti-Dig Fab (Roche Diagnostics, Mannheim, Germany).

Slc30a8_f: 5’-TCAGTCTGTGTTCGCTCTGG-3’,

Slc30a8_r: 5’-TTTCTCGAAGCACCTCCTGT-3’,

Mt2_f: 5’-ATTTCTAAGGAACTTTCAAGC-3’,

Mt2_r: 5’-TTACAGACATACGATTTAGGTGACACT-3’,

Tyr_f: 5’-TTACAACCAAACCTGCCAGTGC-3’,

Tyr_r: 5’-ACTGAAGACATGGAGCCGTTCA-3’.

### Whole mount histochemistry

Dithizone (Sigma-Aldrich, St. Louis, Missouri) (20 mg/mL) was dissolved in DMSO with 0.1 M Tris base (Yuan, 2011) and used 1:1000 in E3. For the von Kossa stain, embryos were incubated with 10% silver nitrate in water under bright light for 10 minutes. For TSQ fluorescence, TSQ (6-methoxy-8-p-toluenesulfonamido-quinoline, Sigma-Aldrich) was dissolved in DMSO to 2 mg/mL and used 1:200 in E3. Larvae were stained for at least one hour and imaged directly with an Olympus SZX16 stereomicroscope (dithizone) or Leica SPE confocal (TSQ).

### Metal determination

Larvae (4 dpf) were anaesthetized and selected based on pigmentation phenotype. Groups of 10 larvae were collected in 2 mL tubes, rinsed with MilliQ water and dried in a vacuum at 60°C. Samples were digested in 1 mL of 3% HNO_3_ overnight at 70°C. Determination was performed by ICP-AES at the department of Earth Sciences, UCL.

### Metallothionein:GFP construct

The 1428-bp metallothionein promoter was cloned with the following primers into a vector containing EGFP and Tol2 sites for stable transgenesis (Kawakami, 2004):

Mt_promoter_f: 5’-AGAGACACTGCACACGTTAC-3’,

Mt_promoter_r: 5’-CAGAGAGTATCCACAA-3’.

Injected larvae were sorted based on GFP expression, raised to adulthood, and outcrossed to establish a stable line.

### Rhodopsin qPCR

Whole-tissue RNA was extracted from larvae at 6 dpf, reverse transcribed with Superscript III, and amplified with SYBR Green qPCR master mix (both from Thermo Fisher, Waltham, Massachusetts) using the following primers:

Rho_qpcr_f: 5’-ACTTCCGTTTCGGGGAGAAC-3’,

Rho_qpcr_r: 5’-GAAGGACTCGTTGTTGACAC-3’.

### Visual function

Electroretinograms were recorded on isolated eyes from 6 dpf zebrafish larvae as previously described (Zang et al., 2015). Briefly, siblings and mutants were dark adapted for half an hour in Ringer’s solution (111 mM NaCl, 2.5 mM KCl, 1 mM CaCl_2_, 1.6 mM MgCl_2_, 10 µm EDTA, 10 mM glucose, and 3 mM HEPES buffer, adjusted to pH 7.7–7.8 with NaOH). Afterwards larvae were placed in the centre of the recording chamber which was filled with 1% agarose. Eyes were removed by pulling the body with forceps while cutting the optic nerve by a Tungsten wire loop. The eye was then repositioned to allow the cornea to face the light source (ZEISS XBO 75W). The recording pipette (1.0 mm O.D. *0.58 mm I.D., GC100F-10, Harvard Apparatus, <u>Holliston, Massachusetts</u>) with a diameter around 20 μm was filled with Ringer’s solution and positioned on the centre of the cornea. 100% light intensity was 591 lux, flash duration was 100 ms with stimulus intervals of 10 s. The stimulus started from 100% intensity and decreased to 0.01% and then went up again to 100%. The b-wave amplitude was calculated as the average of two responses.

The optokinetic response (OKR) was performed as described (Huber-Reggi et al., 2012; Rinner et al., 2005). The larva was immobilized by being embedded dorsal-up in a 35 mm petri dish filled with prewarmed (28°C) 3% methylcellulose. In this condition the eyes can move freely while body movements are restrained. The petri dish was then placed in the centre of a drum with black and white gratings projected by computer generated stimulus via an LCD projector (PLV-Z3000; Sanyo). Both eyes were stimulated at a maximal illumination of 400 lux. To determine contrast sensitivity, a spatial frequency of 20 cycles/360° and an angular velocity of 7.5 deg/s was used with varying contrast (5%, 10%, 20%, 40%, 70% and 100%). To determine spatial sensitivity, an angular velocity of 7.5/s and 70% of the maximum illumination was used with varying spatial frequency (7, 14, 21, 28, 42, 56 cycles/360°). To determine temporal sensitivity, maximum illumination and 20 cycles/360° were used with varying temporal frequency (5, 10, 15, 20, 25, 30 deg/s).

Statistical analysis was performed by SPSS (IBM) using repeated-measures ANOVA. The ERG graphs were generated by Excel and OKR graphs were generated by SPSS.

### Retinal histology

Fixed larvae were embedded in JB-4 resin (Sigma Aldrich) and 10 µm sections were cut with a microtome. Sections were stained with BODIPY TR methyl ester (Thermo Fisher) diluted 1:200, SYTOX Green (Thermo Fisher) 1:30000, and/or TSQ 1:200, all for 30 minutes. BODIPY foci were counted along the retinal OS layer and normalised for OS length.

## Acknowledgements

S.S.-M. acknowledges support from the Smart Mix Programme of the Netherlands Ministry of Economic Affairs, the European Space Agency, and TreatOA (Translational Research in Europe Applied Technologies for OsteoArthritis, FP7). The authors wish to thank S. Carter, L. Lleras, I. Bianco, T. Hawkins and K. Tuschl for valuable advice and manuscript proofreading. This study was supported by a Wellcome Trust Investigator Award (095722/Z/11/Z) to SW and an MRC Programme Grant (MR/L003775/1) to SW and Gaia Gestri.

## Author contributions

E.M. performed experiments and analysed data, except for visual function studies and analysis (J.Z. and S.N). E.M., S.N., S.S.-M. and S.W. conceived experiments and E.M wrote the manuscript with input from S.N, S.W and S.S.-M. J.P.M contributed to positional cloning of the *hu1798* allele. S.I.M contributed eye histology work and L.H.T contributed ISH work.

---

**Figure S1:**
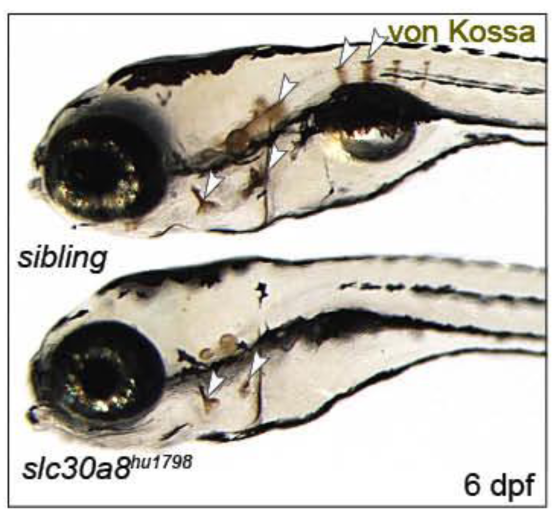
The von Kossa stain shows a reduction of phosphate in *slc30a8*^*hu1798*^ larvae. Arrowheads indicate bone elements.

**Figure S2:**
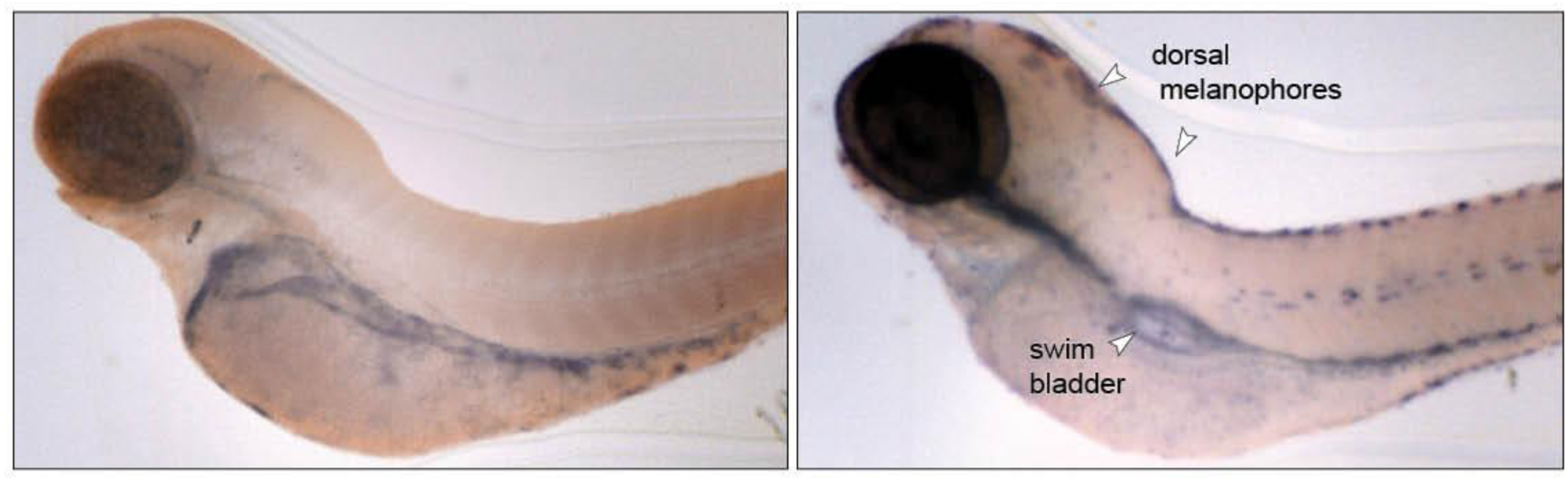
comparison of ISH staining using NBT-BCIP (blue) for *slc30a8* (left) and *tyr* (tyrosinase, right). Unlike *slc30a8, tyr* is detected in pigmented cells (arrowheads). The larvae here are 4 dpf *slc30a8*^*hu1798*^ *a*nd were bleached with peroxide prior to staining to remove endogenous pigment.

**Figure S3:**
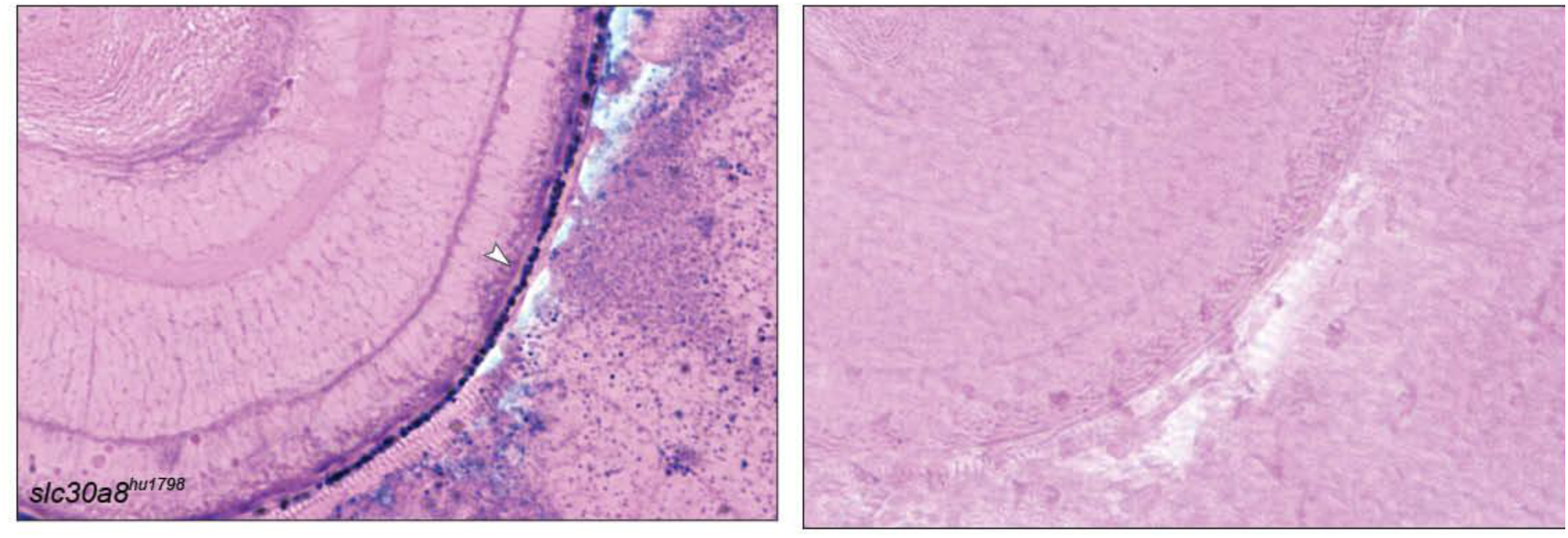
A section of an *slc30a8*^*hu1798*^ eye labelled with an *in situ* hybridization probe for *slc30a8*, demonstrating expression in the RPE. Arrowhead indicates a pattern of *slc30a8* expression (blue NBT-BCIP). Pink counterstain: eosin. Right: a sibling eye section stained with the same probe. Both sections were bleached to remove endogenous pigment.

